# A peripherally restricted cannabinoid 1 receptor agonist provides analgesic benefit from neuropathic pain and a lack of addiction-related behavior

**DOI:** 10.64898/2026.04.13.718281

**Authors:** Amie Severino, Lindsay M. Lueptow, Emily Ellis, Dania Alkoraishi, Igor Spigelman, Catherine M. Cahill

## Abstract

**Introduction:** Cannabis is increasingly used for pain management, with many patients reporting relief from chronic pain that did not respond to conventional treatments. However, cannabis is also associated with unwanted side effects including psychomimetic effects and the potential of developing a cannabis use disorder. To circumvent the central nervous system effects, we investigated whether a peripherally restricted cannabinoid receptor (CB1) agonist, PrNMI [(4-{2-[-(1E)-1[(4-propylnaphthalen-1-yl)methylidene]-1H-inden-3yl]ethyl}morpholine] attenuated pain hypersensitivity associated with nerve injury and profiled its’ abuse potential.

**Materials and Methods:** Mice with chronic constriction injury (CCI) of the sciatic nerve developed hypersensitivity to mechanical stimulation. Paw withdrawal thresholds were assessed following administration of PrNMI (i.p. 0.3 mg/kg and 0.6 mg/kg) or vehicle in CCI and sham mice. The conditioned place preference model was used to measure drug-reward to 0.6 mg/kg i.p. PrNMI in CCI and sham-injury control animals. We further assessed abuse potential to determine if PrNMI (0.5 mg/kg) would reinstate drug-seeking behavior in mice trained to self-administer intravenous fentanyl (10 μg/kg/infusion).

**Results:** PrNMI administration transiently increased paw withdrawal thresholds in mice with CCI-induced allodynia in a dose-dependent manner. PrNMI conditioning did not produce a conditioned place preference in mice with either CCI or sham injury. Mice who had learned to self-administer fentanyl and went through extinction training did not reinstate drug-seeking behavior when administered PrNMI.

**Discussion:** The systemic CB1 receptor agonist PrNMI demonstrated analgesic benefit in alleviating mechanical allodynia associated with chronic constriction injury of the sciatic nerve without increasing addiction related behaviors associated with the establishment of addiction.

## Introduction

Self-medication of cannabinoid drugs including cannabis and derived formulations is highly prevalent, including its use for pain management. Adults with chronic pain conditions frequently use medicinal cannabis to treat their pain, and some report that the use of cannabis allows them to reduce their reliance on opioids for pain management (Bicket et al., 2023; National Academies of Sciences, Engineering, and Medicine; Health and Medicine Division; Board on Population Health and Public Health Practice, 2017). Medicinal cannabis users most frequently cite chronic pain as their motivation for obtaining a license to use cannabis, followed by other purported benefits in treating multiple sclerosis symptoms, and nausea and vomiting associated with chemotherapeutics (Boehnke et al., 2022). However, up to 25% of chronic pain patients using cannabis develop clinically diagnosed cannabis use disorder (Dawson et al., 2024). Additionally, the FDA has not approved medicinal cannabis, its’ derived formulations, or synthetic cannabinoids for pain management. Strategies are needed to treat pain with little to no abuse liability or the potential for addiction, including selective targeting of peripheral cannabinoid receptors.

Chronic pain is characterized by lasting hypersensitivity to previously non-painful stimuli, a phenomenon called allodynia. Allodynia can be in response to thermal or mechanical touch to the skin, which often persists beyond the initial healing of the external injury, surgery, or cessation of neuropathy inducing drug exposure (Glare et al., 2019; Melzack and Wall, 1965; Mulpuri et al., 2018; Sandkühler, 2009; AL Severino, Chen, et al., 2018). In humans, chronic pain increases the likelihood of exposure to prescription opioids and the subsequent potential for misuse (Cahill et al., 2013; Chang and Compton, 2013; Nazarian et al., 2021; Severino, Shadfar, et al., 2018; Volkow and McLellan, 2016). The long term use of opioid medications can also create a tolerance to analgesic benefits and even increased pain, a phenomenon called opioid-induced hyperalgesia (Mercadante et al., 2019; Von Korff et al., 2011). This suggests that eventually patients might need to change medications for relief.

Cannabinoid drugs could be an alternative strategy to reduce prolonged opioid exposure to decrease the potential for developing addiction. Evidence from the cannabis usage patterns and off label indications for cannabinoid drugs suggests that cannabinoids can reduce pain (Anthony et al., 2020; Bicket et al., 2023; Boehnke et al., 2022; Gregus and Buczynski, 2020; Jergova et al., 2021; Lowin and Straub, 2015; Milligan et al., 2020; Mulpuri et al., 2018; Rabgay et al., 2020; Reiman et al., 2017; Villanueva et al., 2022; Wang et al., 2019; Woodhams et al., 2017). The viability of cannabinoids as analgesics rests upon the ability to reduce the adverse side effects of cannabis (Blume et al., 2016; Hoch et al., 2025; Hryhorowicz et al., 2019; Ignatowska-Jankowska et al., 2015; Mulpuri et al., 2018; Shen et al., 2024; Tian et al., 2025; Wouters et al., 2019). Cannabis and its synthetic or derived compounds demonstrate potential negative side effects that are mediated by binding to cannabinoid receptors in the central nervous system (Hoch et al., 2025; Mackie, 2005; Woodhams et al., 2017). Adverse side effects of cannabis include intoxication, gastrointestinal effects, cardiovascular effects, and potential for developing Cannabis Use Disorder (Hoch et al., 2025). The cannabinoid system has a demonstrated interaction with the opioid receptor system. Cannabinoid analgesics should be evaluated for cross-addictive potential when considering that many patients with chronic pain are likely to use prescription opioids for pain management.

One strategy for reducing cannabinoid side effects is to selectively target peripheral cannabinoid receptors that are localized on peripheral nociceptor sensory neurons. This strategy would bypass the central nervous system effects of cannabinoids and is a viable analgesic target (Volkow and McLellan, 2016; Wiese et al., 2022). It has been previously demonstrated that peripheral cannabinoid receptor 1 (CB1R) on voltage gated sodium channel 1.8 expressing nociceptive receptors mediate the analgesic properties of cannabinoids (Agarwal et al., 2007). Thus, the nociceptor-specific loss of CB1 substantially reduced anti-hyperalgesic effects produced by local and systemic, but not intrathecal, delivery of cannabinoids in models of chronic inflammatory and neuropathic pain. The peripheral cannabinoid receptor agonist [(4-{2-[-(1E)-1[(4-propylnaphthalen-1-yl)methylidene]-1H-inden-3yl]ethyl}morpholine] PrNMI is an indole cannabimimetic compound that demonstrates analgesic potential in a chemotherapy-induced model of neuropathy and cancer induced bone pain, without significant central nervous system mediated side-effects (Mulpuri et al., 2018; Seltzman et al., 2016; Zhang et al., 2018). Here we demonstrated that PrNMI provided analgesic benefit in the chronic constriction injury model of neuropathic pain that did not cause addiction related behaviors associated with either reward using the conditioned place preference model or cross-addictive potential in opioid self-administration reinstatement in mice.

## Methods

### Animals

All experiments used male and female C57BL/6J mice between 8-12 weeks age (The Jackson Laboratory, Bar Harbor, ME). Unless otherwise stated, mice were group housed, fed standard mouse chow and water *ad libitum* with exceptions for the durations of experimental testing. Mice were maintained in a temperature-controlled vivarium on a reverse light-dark cycle (lights off between 8 am and 8 pm). All procedures were conducted in accordance with the National Institutes of Health Guide for the Care and Use of Laboratory Animals, were preapproved by the University of California Los Angeles Institutional Animal Care and Use Committee, and were compliant with ARRIVE 2.0 guidelines (*Guide for the Care and Use of Laboratory Animals*, 2011; Percie du Sert et al., 2020). After a week of vivarium acclimatization, all mice were handled by experimenters and acclimated to experimental apparatuses for at least 1 week prior to surgical, drug, or behavioral interventions. Mice were habituated to the behavior room in their home cages for 10-20 minutes before being handled each day. Estrous cycle phases were not tracked in female mice because of the sex-specific additional stress and that biological sex was not a primary experimental outcome. Both sexes were used in these studies and tested at separate times, where the opposite sex was not in the behavioral testing room. All equipment was thoroughly cleaned between testing male and female mice.

Mice were assigned to conditions in a randomized block design so that the running of subjects counterbalanced factors such as time of day over experimental conditions. We completed experiments in successive replications where replications will be balanced with respect to experimental groups. All behavior was performed in the dark (active) phase of the reverse light-dark cycle and recorded by infrared camera using an infrared illuminator under low lighting conditions (<10 LUX).

Separate groups of mice were used for each of the behavioral outcomes where a total of 79 mice used for all studies. Mechanical paw withdrawal threshold testing was conducted in equal numbers of male and female mice divided by surgical/pain condition (n= 8 per sex per surgical condition) for a total of 32 mice. Conditioned place preference experiments were conducted on equal numbers of male and female mice (n= 8 per sex per surgery condition) for a total of 32 mice. Intravenous self-administration experiments were conducted on male pain naïve mice (n= 15).

## Surgical procedures

### Chronic constriction injury (CCI)

Chronic constriction injury of the left sciatic nerve or sham surgeries were conducted in mice as we previously reported in order to produce hypersensitivity to touch that can be attenuated by analgesic medications (Liu et al., 2019; Lueptow et al., 2025). Temperature conditions were maintained throughout procedures using a heating pad and eyes were treated with lubricating ointment to prevent drying. Mice were anesthetized with isoflurane (5% induction, 2% maintenance) and the surgical site was prepared by shaving and 3 alternating swabs of alcohol and betadine. A superficial incision was made, the biceps femoris muscle was resected, and a 2-mm cuff constructed of PE20 polyethylene tubing (Intramedic) was placed on the main branch of the left sciatic nerve.

The sham condition mice were induced and maintained on isoflurane anesthesia for a similar period of time as the experimental group mice. In sham mice, the surgical site was shaved and sterilized, but no incision was made. All mice received acetaminophen (∼3 mg, p.o.) following surgery as part of postoperative care and again 24 hours later. Mice also received a subcutaneous injection of 1 mL of lactated Ringer solution after surgery to maintain hydration. All mice were allowed to recover from surgery for a minimum of 96 hours before further handling and experimentation. Sham surgery consisted of similar duration of anesthesia, skin incision and post operative treatment, but no dissection of the nerve.

### Jugular catheterization

Mice were implanted under aseptic conditions with catheters in the right jugular vein as previously described (Hakimian et al., 2019; Lueptow et al., 2025). Isoflurane anesthesia (5% induction and 2% maintenance) was used. In brief, 12 mm of 1 Fr silicone tubing (0.2 mm i.d., 0.4 mm o.d., Norfolk Access, Skokie, IL) was inserted into the right vein and connected to a back-mounted 26-gauge stainless-steel guide cannula (315BM-8-5UP, Plastics One, Roanoke, VA). Catheters were flushed daily with 0.02 to 0.04 mL of sterile saline mixed with heparin (30 U/mL) and cefazolin (10 mg/0.1 mL). Two days after surgery, catheter patency was tested with an infusion of 0.02 mL of propofol (10 mg/mL) through the jugular catheter and at the end of experimentation. The surgery success rate was ∼75% successful in obtaining patent cannulated mice. Only mice that remained patent to the end of the progressive ratio task were included in the study.

## Behavioral Testing

### Mechanical Withdrawal Threshold

To assess the dose response and the time course of PrNMI analgesia, 50% paw withdrawal thresholds were assessed at baseline and 1 hr, 1.5 hr, 2 hr, and 24 hr following subcutaneous administration of either saline, 0.3 mg/kg PrNMI or 0.6 mg/kg PrNMI at 13-14 days post-CCI surgery. The 50% paw withdrawal threshold was measured using von Frey filaments in a modified up-down method (Bonin et al., 2014; Chaplan et al., 1994; Liu et al., 2019; A Severino, Chen, et al., 2018). Mice were habituated to the von Frey apparatus for 15 minutes prior to experimentation by being placed in individual plexiglass chambers on top of a mesh floor. Each von Frey filament (Stoelting, Wood Dale, IL) was applied to the paw for 5 seconds to determine if a response was elicited. If a response was elicited, the force was increased for the next filament. If a response was not demonstrated, the force was decreased for the next filament. This continued for 8 trials and the 50% paw withdrawal threshold was calculated (Bonin et al., 2014). A positive withdrawal response was defined as lifting or moving the paw away from the filament except where it was ambulation.

### Conditioned place preference

This test was conducted using an unbiased, counter-balanced three chamber apparatus as previously described (Cahill et al., 2013; Liu et al., 2019; Taylor et al., 2015). Each box (28 × 28 × 19 cm) was divided into two equal-sized conditioning chambers and a neutral compartment. The two chambers were distinguished with visual, scent, and tactile cues. Mice were placed in the apparatus and allowed free access to both chambers for habituation to the apparatus for 15 minutes. The following day, animals were assessed for pre-conditioning bias to conditioning chambers prior to drug pairing assignment. Mice were placed in the apparatus with free access to all chambers for 30 minutes with ANY-maze video tracking to determine time spent in each of the chambers. Animals then underwent either CCI surgery or sham surgery as described above. On day 8 post-surgery, mice were conditioned to subcutaneous PrNMI (0.6 mg/kg) or vehicle and isolated to the paired chamber for 30 minutes. Mice were conditioned once a day to either vehicle or PrNMI for an additional 5 days for a total of 3 drug and 3 vehicle pairings. Day, drug and chamber were counterbalanced. Post-conditioning (drug-free test) was performed the day following the last conditioning session where animals were again allowed to explore the entire apparatus for 30 minutes. Time spent and activity in each compartment were recorded during the sessions with ANY-maze software. A state-dependent test was performed two days after the post condition test where all mice were injected with the same dose of PrNMI used for conditioning. Preference scores were calculated using the following formula: [Time in Test(paired) − Time in Test(unpaired)] − [Time at baseline(paired) − time at baseline(unpaired)].

### Operant intravenous self-administration of fentanyl

Experiments were conducted in 8 operant conditioning chambers (Med Associates, St. Albans, VT), as previously described (Hakimian et al., 2019; Lueptow et al., 2025). Each conditioning chamber was fitted with 5-unit nose poke ports, house and cue lights (ENV-115C, Med Associates, Fairfax, VT), and a multiaxis lever arm (PHM-124 MW, Med Associates) connected to a syringe pump (PHM-100, Med Associates, Fairfax, VT) in sound-attenuating boxes with ventilation fans. The cannulas for drug delivery were connected to PE20 tubing that was threaded into the multiaxis lever arm.

Two nose-poke ports were available in each operant box, 1 designated *active* to deliver drug and 1 *inactive* where there was no consequence. Cue lights were on above both active and inactive ports. Upon a nose poke into the active port, the light was turned off, the drug was delivered for 2 seconds, and the drug became unavailable for an additional 18 seconds for a total of 20 seconds time-out (TO). Additional nose pokes made during the time-out were recorded. Inactive nose pokes were recorded but resulted in no changes to cues or time out. Active and inactive nose-poke assignments were counterbalanced on the left and right sides of the operant boxes.

One week after the jugular vein catheterization, all mice were trained to self-administer fentanyl in a 6 hour autoshaping session (Spectrum Chemicals, 10 μg/kg/infusion, prepared in sterile saline) under a fixed ratio 1 (FR1) schedule, where mice received 0.67 μL/g body weight/2 seconds. Fentanyl was chosen for this study because it is a commonly diverted opioid drug with fast onset of action and short duration of effect, allowing animals to learn operant behavior quickly(Chen et al., 2025; Comer and Cahill, 2019). During this autoshaping day, active and inactive ports were baited every 15-30 minutes with Ensure. The day following autoshaping, mice had access to the boxes for 2h or received 100 drug infusions (whichever came first) on an FR1 schedule with a 20 second time out. Mice were tested 5 days a week with no behavioral testing on weekends. There was no significant effect of the drug holiday on subsequent drug taking (data not shown).

Mice maintained self-administration behavior for 10 days on FR1 schedule prior to extinction training for an additional 10 days, where no cues or drug were administered. Extinction was defined as animals not exhibiting a preference for the active port as well as making less than 10 active port pokes. Following the extinction phase, a reinstatement session was performed on three separate days. On the first day of reinstatement test, all mice received vehicle injections. On the second day of the reinstatement test, all mice received PrNMI (0.5 mg/kg) injections. Both active and inactive nose pokes were determined over a 2h period for each day. Fentanyl (10 μg/kg) was administered on the final day of reinstatement testing to assess drug-priming reinstatement of drug-seeking in mice that did not show any evidence of PrNMI reinstatement (no difference from their last day of extinction training) in a 1 hour session.

### Statistical analysis

Experiments were designed to ask (1) does PrNMI attenuate pain hypersensitivity in a model of chronic neuropathic pain, (2) does PrNMI elicit either positive or negative reinforcement (preference in a chronic pain state) and (3) does PrNMI induce reinstatement of drug-seeking behavior. All data sets were tested for normality and *post hoc* tests were selected based on the results. All statistical analysis was conducted on GraphPad Prism (v10). Mechanical withdrawal thresholds were analyzed by a repeated measures 2-way ANOVA followed by Sidak’s multiple comparison *post hoc* analysis. Conditioned place preference data was analyzed with raw data as time in chamber using a 2-way ANOVA followed by a Sidak’s multiple comparison *post hoc* analysis. The conditioned place preference score was analyzed by an unpaired t-test as well as a one sample t-test to determine if scores were significantly different than a theoretical value of zero (no preference/aversion). Self-administration time course data were analyzed by using a repeated measures 2-way ANOVA. All time course data were performed by repeated measures ANOVAs using a Geisser–Greenhouse correction. Drug reinstatement testing was analyzed by a one-way ANOVA followed by Tukey’s multiple comparison test.

Significance was denoted as p<0.05.

## Results

### Pain was transiently attenuated in a dose-dependent manner by peripheral CB1R agonist PrNMI

The effects of PrNMI were determined 2 weeks post injury (Figure 1A). In animals with chronic pain, PrNMI treatment was associated with reversible increases in 50% paw withdrawal thresholds on the paw ipsilateral to the site of CCI in both males and females (Figure 1B). A two-way ANOVA revealed a significant effect of time (F_(2.373,106.8)_ = 64.24, ****p<0.0001), drug treatment (F_(2,45)_ = 83.98, ****p<0.0001) and an interaction (F_(4.746,106.8)_ = 17.57, ****p<0.0001). There was a main effect of PrNMI treatment over time in the ipsilateral side to the injury that also occurred in the paw contralateral to the nerve injury (Figure 1C), but only at the highest dose. A two-way ANOVA revealed a significant effect of time (F_(3.526,158.7)_= 3.790, **p=0.0080), drug treatment (F_(2,45)_= 8.694, ***p=0.0006) and an interaction (F_(7.052,158.7)_= 4.057, ***p=0.0004). The peak of anti-allodynic effect was at 2 hours and paw withdrawal thresholds returned to baseline pain levels after 24 hours. Data are also presented by separating sexes (Figure 1B,C). There were no sex differences whereby both females and males exhibited PrNMI-induced attenuation of mechanical hypersensitivity. In female mice, a two-way repeated measures ANOVA revealed a significant main effect of treatment (F_(2,21)_ = 27.00, ****p<0.0001), a main effect of time (F_(3.394,71.27)_ = 85.49, ****p<0.0001), as well as a time x treatment interaction (F_(3.394,71.27)_= 11.26, ****p<0.0001) in the ipsilateral side to injury. For the contralateral hind paw, a two-way repeated measures ANOVA revealed a significant main effect of treatment (F_(2,21)_ = 5.214, *p=0.0145), a main effect of time (F_(3.689,77.48)_ = 2.804, *p=0.0351), as well as a time x treatment interaction (F_(7.379,77.48)_ = 2.804, **p=0.0096). In male mice, a two-way repeated measures ANOVA revealed a significant main effect of treatment (F_(2,21)_ = 39.64, ****p<0.0001), a main effect of time (F_(2.712,56.95)_ = 49.37, ****p<0.0001), as well as a time x treatment interaction (F_(5.424,56.95)_= 7.342, ****p<0.0001) in the ipsilateral side to injury. For the contralateral hind paw, a two-way repeated measures ANOVA revealed a significant main effect of treatment (F_(2,21)_ = 11.82, ***p=0.0004), a main effect of time (F_(3.454,72.54)_ = 5.340, **p=0.0014), as well as a time x treatment interaction (F_(6.909,72.54)_= 6.725, ****p<0.0001).

**Figure 1.**
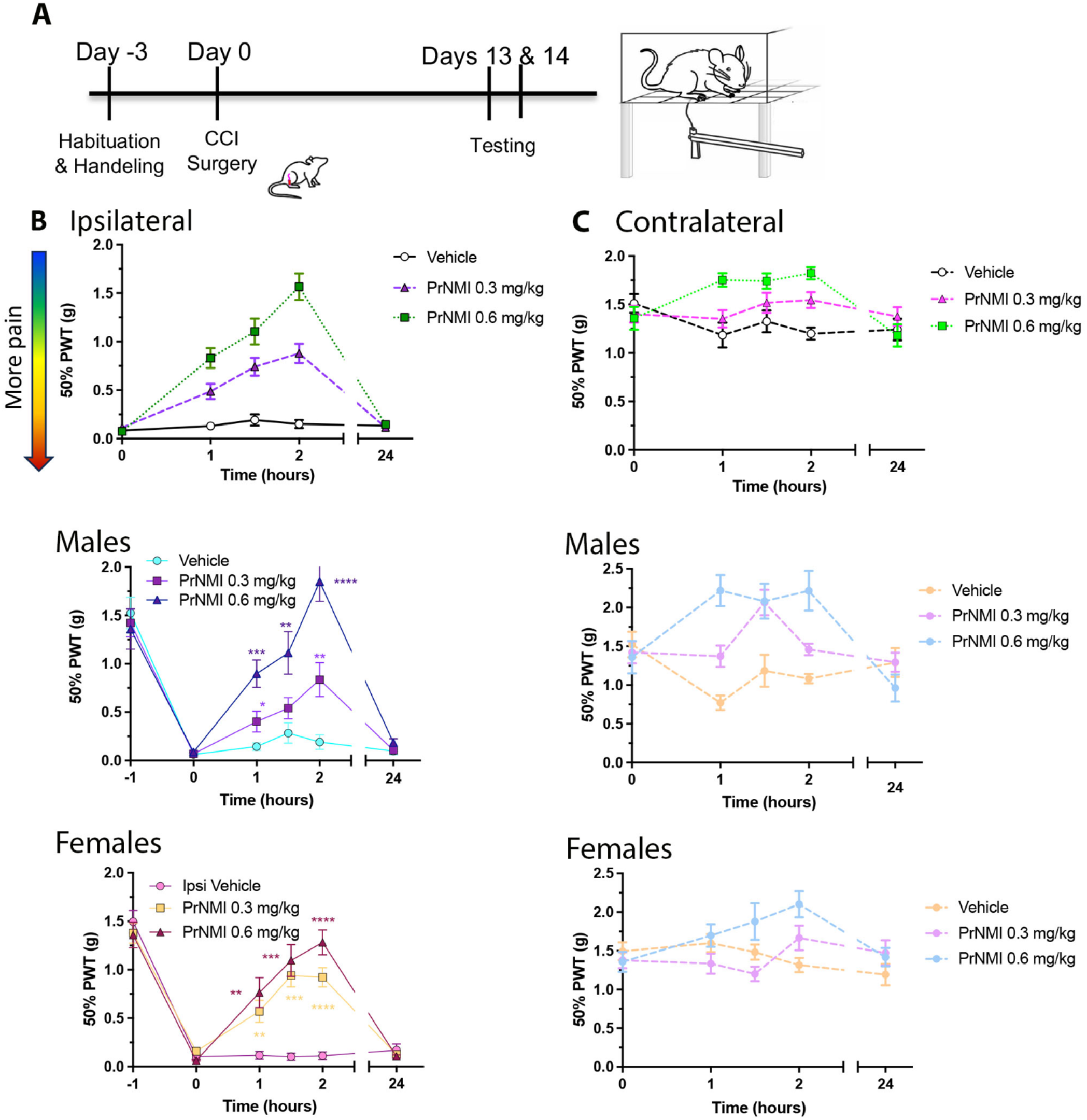
Systemic administration of PrMNI attenuated mechanical hypersensitivity associated with sciatic nerve injury in male and female mice. A. Schematic of experimental timeline. B. PrMNI dose-dependently increased von frey mechanical withdrawal thresholds compared to vehicle treatment ipsilateral to nerve injury in a model of neuropathic pain. Top panel represents the combination of male and female data, whereas in the lower panels, data was divided by sex (N= 16 per group). C. PrMNI increased von frey mechanical withdrawal thresholds compared to vehicle treatment on the contralateral side in a model of neuropathic pain. Top panel represents the combination of male and female data, whereas in the lower panels, data was divided by sex (N= 16 per group). Data was analyzed by a repeated measures 2-way ANOVA and presented as means +/-S.E.M.

### Peripheral CB1R agonist PrNMI did not produce reward behavior

The timeline of the experimental protocol to determine the potential rewarding effects of the CB1R agonist PrNMI is presented in Figure 2A. The administration of PrNMI did not elicit a conditioned place preference in animals with or without CCI pain (Figure 2B-E). Post-condition testing in the absence of drug showed no difference between the time spent in the vehicle vs drug chamber for either sham or CCI mice (Figure 2B). A two-way ANOVA revealed no significant effect of treatment (F_(1,58)_= 0.00091, p=0.9242), pain condition (F_(1,58)_= 0.0315, p=0.8597), nor an interaction (F_(1,58)_= 1.601, p=0.2108). These data were confirmed by calculating a conditioned place preference score that takes into account pre-conditioning bias (Figure 2C), where an unpaired Welsh’s t-test showed no significant difference between groups (t=1.391, df=28, p=0.1748). A one-sample t-test also showed that no group was significantly different than a theoretical value of zero, demonstrating that the drug did not produce either a preference or aversion. Two days following the post-condition test, all mice were injected with the PrNMI to test the state-dependent preference (Figure 2D). A two-way ANOVA revealed no significant effect of treatment (F_(1,58)_ = 0.1218, p=0.7283), pain condition (F_(1,58)_ = 0.0211, p=0.8850), but did show an interaction (F_(1,58)_= 4.742, p=0.0335). Analysis of the conditioned place preference score (Figure 2E) showed no difference between sham and CCI mice (t=1.002, df=28, p=0.3250), and the one sample t-test also did not identify any effect.

**Figure 2.**
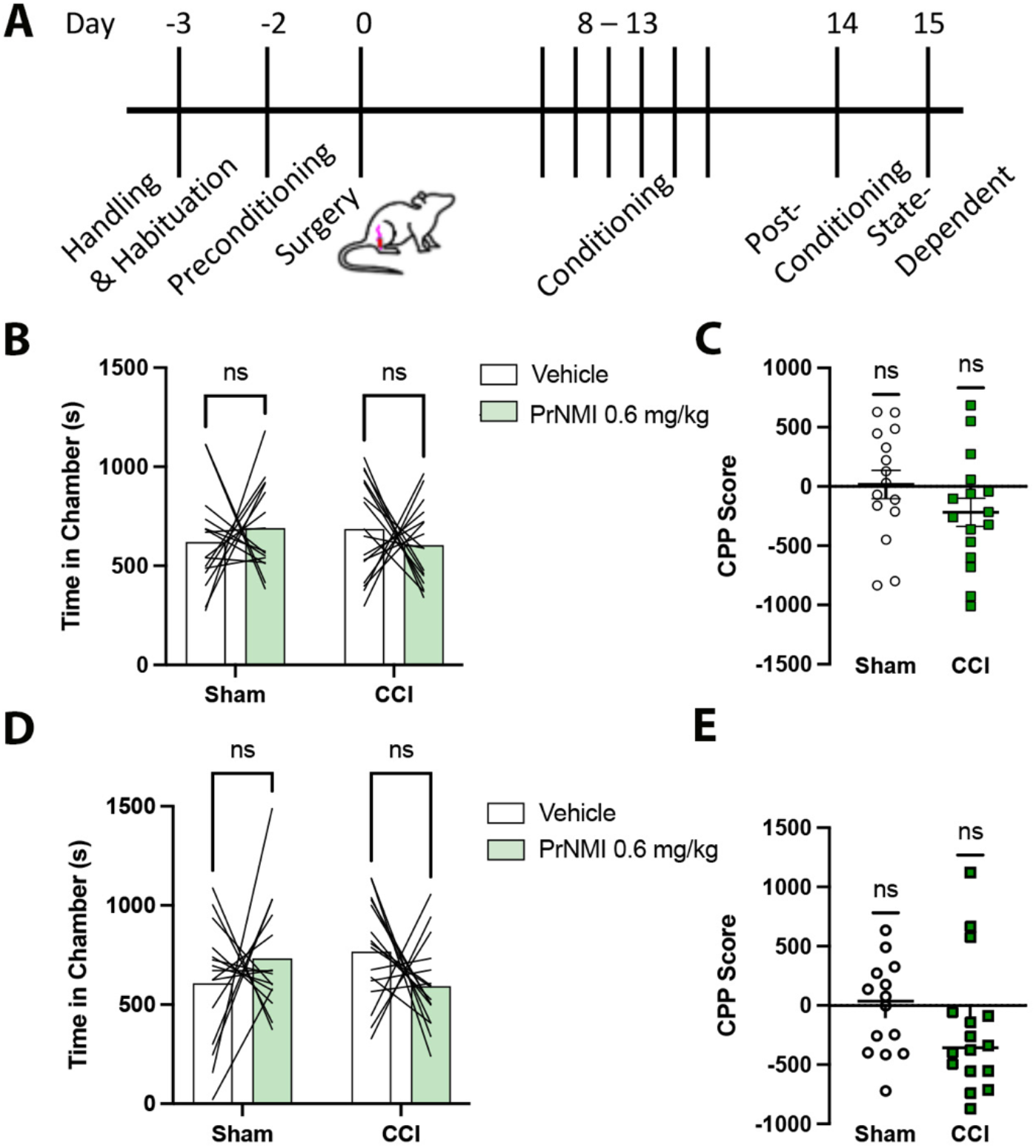
PrMNI did not produce reward in either pain-naïve or chronic neuropathic pain animals. A. Schematic of experimental timeline. Mice were conditioned once a day for 6 days counterbalanced for treatment day and contextual cues prior to testing in a drug free state. B. Post condition test in the absence of drug demonstrates that PrMNI did not increase time spent in the drug-paired chamber. C. A conditioned preference score (CPP) was calculated by taking into account any potential initial contextual bias. There was no effect of the PrMNI in either sham or CCI mice as evidenced by no difference between groups, and no difference in a one sample t-test by comparing to a theoretical value of zero (N=16 per group). D. State-dependent test in the presence of drug demonstrates that PrMNI did not increase time spent in the drug-paired chamber. C. A conditioned preference score (CPP) demonstrated that there was no effect of the PrMNI in either sham or CCI mice as evidenced by no difference between groups, and no difference in a one sample t-test by comparing to a theoretical value of zero (N=16 per group). Data represent means +/-S.E.M.

### Peripheral CB1R agonist PrNMI did not produce reinstatement behavior in the IVSA model of fentanyl addiction

The timeline of the experimental protocol to determine the potential of the CB1R agonist PrNMI to re-instate drug seeking behavior (Figure 3A). Animals acquired fentanyl IVSA that was maintained by increased nose pokes in the active port paired to fentanyl administration (Figure 3B) where a repeated measures 2-way ANOVA revealed a significant effect of time (F_(4.112,112.4)_= 3.034, *p=0.0194) and nose poke (F_(1,28)_= 21.43, ****p<0.0001), but not interaction (F_(9,246)_ = 1.183, p=0.3060). Following 10 days of self-administration, mice underwent extinction behavior where no drug or cues were available to the mice. Compared to the last day of when the fentanyl was available, mice drug-seeking behavior evidenced by an increase in both active and inactive nose pokes (Figure 3C). A repeated measures ANOVA revealed a significant effect of time (F_(3.103,85.63)_ = 14.18, ****p<0.0001) and nose poke (F_(1,28)_ = 4.219, *p=0.0494), but not interaction (F_(10,276)_ = 0.9257, p=0.5098). The average time to extinction was 9 days post-removal of fentanyl. To determine the extent the PrNMI drug could reinstate drug seeking behavior for the fentanyl, we administered vehicle or PrNMI to all animals in a randomized cross over design. PrNMI did not produce an increase in active nose pokes (Figure 3D) or total nose pokes (Figure 3E) compared to vehicle treatment. Following the PrNMI all mice were injected with fentanyl as a positive control to induce reinstatement. Fentanyl in the absence of cues showed reinstatement for drug seeking as evidenced by an increase in active nose pokes (Figure 3D) and total number of nose pokes (Figure 3E). A one-way ANOVA of active pose pokes revealed a significant effect of treatment (F_(3,41)_ = 5.027, **p=0.0047). Tukey’s multiple comparison *post hoc* analysis showed that only the fentanyl caused reinstatement. Similar statistics were identified for total nose pokes where a one-way ANOVA of active pose pokes revealed a significant effect of treatment (F_(3,44)_= 5.789, **p=0.0020). Tukey’s multiple comparison *post hoc* analysis showed that only the fentanyl caused reinstatement.

**Figure 3.**
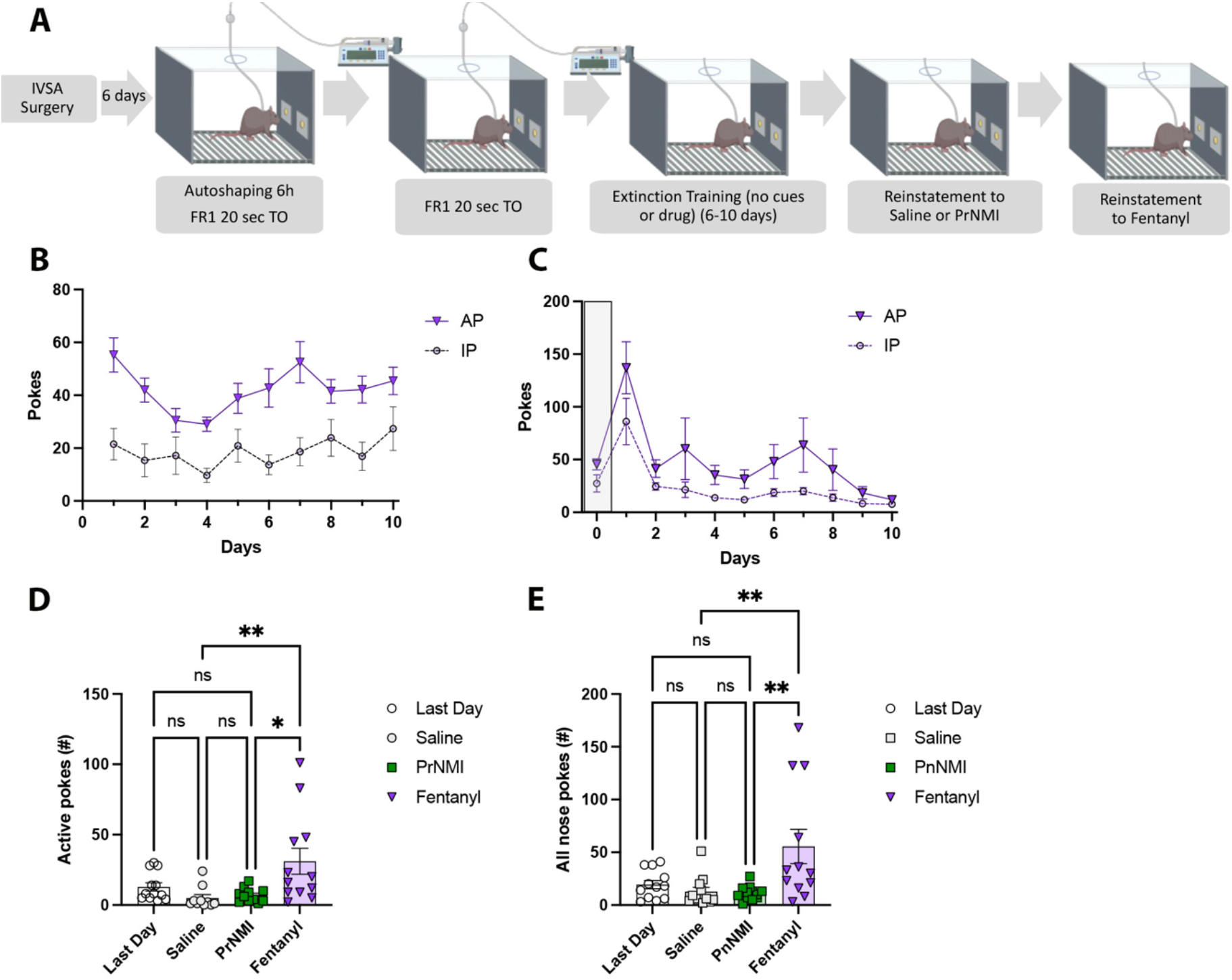
prNMI did not substitute for oxycodone self-administration nor trigger drug relapse following extinction. A. Schematic of experimental timeline. Mice underwent training to self-administer fentanyl (10 ug/kg/infusion for 2h access) prior to extinction training and reinstatement test. B. Active and Inactive nose pokes for fentanyl self-administration. C. Mice show burst of drug-seeking when the fentanyl is removed and all extinguish their drug seeking by 10 days of training. D. After the last day of extinction, mice were counter-balanced for either vehicle (saline) or PrNMI injection. Neither saline or PrNMI induced reinstatement of drug-seeking as assessed by active nose pokes. After the two test days with vehicle or PrNMI, all mice received fentanyl and showed a state-dependent drug induced reinstatement of drug seeking. Data was analyzed by a one-way ANOVA (N=11). E. Neither saline or PrNMI induced reinstatement of drug-seeking as assessed by total number of active and inactive nose pokes. After the two test days with vehicle or PrNMI, all mice received fentanyl and showed a state-dependent drug induced reinstatement of drug seeking. Data was analyzed by a one-way ANOVA (N=11). Data represent mean +/-S.E.M.

## Discussion

We found that systemic peripheral CB1R agonist administration was associated with attenuation of pain hypersensitivity associated with nerve injury in both male and female mice. This was demonstrated by increased paw withdrawal thresholds in mechanical hypersensitivity testing. Animals with hypersensitivity to touch after CCI had a significant transiently increased ability to tolerate mechanical forces to paw on the ipsilateral side to the nerve injury and produces antinociceptive effects to the mechanical stimulation on the contralateral side. This anti-allodynic effect was greater at the 0.6 mg/kg dose compared to a 0.3 mg/kg dose of PrNMI overall. However, PrNMI was not associated with behaviors used to assess reward or the establishment or the relapse phases of substance use disorders.

Animals did not exhibit positive or negative reinforcement learning of drug effects with contextual cues modeled by conditioned place preference in either pain-naïve or pain states, suggesting that PrNMI consistently does not produce reward or shift the typical reward profile during pain. PrNMI did not produce reward in the pain animals, demonstrating a lack of effect on affective dimensions of the pain experience. This is consistent with evidence that peripheral CB1 receptors are not involved in the cannabinoid reward system, and rather that central CB1 receptors mediate cannabinoid reward (Kunos et al., 2009). Central nervous system CB1 receptor expression and G-protein recruitment was increased in an animal model of pain, suggesting that caution should be applied to any CB1 receptor approach for pain therapy to minimize off-target central nervous system effects (Wilson-Poe et al., 2021). This is particularly relevant as CB1Rs in cortical glutamatergic neurons were identified to contribute to the negative affective component of pain (Bajic et al., 2018). Animals did not demonstrate cross-reinstatement to PrNMI after opioid extinction in the intravenous self-administration model of fentanyl addiction. This suggests that peripheral cannabinoids may not be rewarding and that in previously established opioid dependence, cannabinoids would demonstrate a lack of cross-addiction potential.

Currently, there are several medicinal compounds approved by the FDA that emulate the psychoactive and non-psychoactive components in cannabis, delta9-tetrahydrocannabinol (THC) and cannabidiol (CBD), respectively. These medications include Marinol (dronabinol) and Cesamet (nabilone), which are similar to THC, or Sativex (nabixmol) and Epidiolex (cannabidiol) that are related to or derived from CBD (Giacoppo et al., 2017; Gregus and Buczynski, 2020; Markovà et al., 2019; Villanueva et al., 2022). These medications are prescribed to treat chemotherapy induced nausea, AIDS induced anorexia, and certain seizure disorders. However, none of these medications are FDA approved for treating pain, a benefit that many people cite as the reason they use cannabis. Additionally, many of these cannabinoid drugs cause central nervous system mediated side effects. There are many strategies in pharmaceutical development that bypass the side effects associated with cannabis, including biased agonist strategies, selective receptor site targeting approaches, or the use of peripherally restricted drugs (Rangari et al., 2025; Seltzman et al., 2016; Shen et al., 2024; Wouters et al., 2019; Zhang et al., 2018). Here we demonstrate that peripheral compartment activity of CB1R agonist is sufficient for attenuating the sensory component of pain without the potential to produce addiction.

The development of a peripheral cannabinoid drug with pain-relieving effects and a lack of addiction-related behavior is a significant advance in cannabinoid research. PrNMI is a peripherally restricted indole cannabimimetic drug that primarily targets CB1 receptors (Mulpuri et al., 2018; Seltzman et al., 2016). This means that PrNMI is synthetic, is not derived or extracted from cannabis, and it acts as an agonist at CB1 receptors in the peripheral nervous system. Indole cannabimimetic compounds have been previously controversial due to potential adverse side effects, and most of the previous research focused on developing CB2R agonists (Howlett et al., 2021). This was to bypass the side effects of central CB1R agonists. However, PrNMI’s analgesic effects are mediated by selective action at peripheral CB1Rs, and does not cause typical central nervous system mediated side effects of cannabinoids (Mulpuri et al., 2018; Seltzman et al., 2016; Zhang et al., 2018). This is promising in tandem with our findings that PrNMI did not elicit the reward associated with the establishment of addiction or a cross-addiction potential for the often diverted medicinal opioid fentanyl.

Substance use disorders have a characteristic behavioral cycle that begins with an establishment of repeated intoxication of a rewarding substance and is maintained by relapses that often follow the periods of drug abstinence. Binges of intoxication occur in an attempt to achieve a reward associated with the drug, leading to an establishment of increased drug-related cue reactivity (Cahill et al., 2016; Evans and Cahill, 2016; Koob, 2021; Uhl et al., 2019). This was assessed in this study using conditioned place preference behavior. Place preference after conditioning occurs in rodent models of morphine exposure, as an example, but also with repeated exposure to other drugs of abuse such as cocaine and amphetamine (Cahill et al., 2013; Cunningham et al., 2006; Grenier et al., 2022; Hou et al., 2023). It has been observed that pain can alter the place preference for environments associated with CNS-penetrant analgesic drugs such as morphine, so it was imperative to assess whether pain experience would alter the place preference for the peripheral cannabinoid PrNMI (Cahill et al., 2013). Mice with CCI injury or a sham-control injury were repeatedly exposed to PrNMI in a chamber that had contextual scent, spatial, and textural cues (Cahill et al., 2013; Cunningham et al., 2006; Green and Bardo, 2020; Grenier et al., 2022; Hou et al., 2023; White and Carr, 1985). We did not observe a place preference with repeated conditioning to PrNMI exposure in animals with or without CCI, suggesting a lack of reward in the pain or pain-free state.

It is essential to study cannabinoids in the context of late-stage opioid dependence to validate whether they cause cross-addictive potential, as chronic pain patients would be likely to switch from opioid medications to a cannabinoid medication. The rodent intravenous self-administration (IVSA) paradigm is a robust behavioral model where animals are implanted with a jugular catheter and taught to associate a cue, such as a light, with an infusion of drug when they perform an action such as pressing a specific lever or poking in a nose port (Chen et al., 2025; Green and Bardo, 2020; Lueptow et al., 2025; McNamara et al., 2010; Ren and Lotfipour, 2022; Slosky et al., 2022). Certain drugs or conditions can potentiate drug addiction related behaviors in the IVSA paradigm (Honeycutt et al., 2022; Lueptow et al., 2025; Windisch et al., 2021). Animals will maintain acquired drug-seeking behavior in the case of opioids such as fentanyl or oxycodone. To model abstinence, the drug and cue were removed. Animals will initially show a burst of drug seeking but will learn to extinguish drug seeking behaviors (Chen et al., 2025; Lueptow et al., 2025; McNamara et al., 2010). To assess the cross-addictive potential of PrNMI, the drug-paired cue was reintroduced and PrNMI was infused to determine if CB1R agonists produce opioid cross-reinstatement. We did not observe an increased cross-addictive potential as evidenced by a lack of cross-reinstatement to PrNMI self-administration after the extinction of fentanyl self-administration.

We found that the peripherally restricted cannabinoid drug PrNMI produced analgesia without demonstrating reward behaviors in pain or pain-naïve states and without cross-addictive potential to opioid dependence behavior. Evaluating this cross-addictive potential in the context of chronic pain would be an avenue of further research, as it is a limitation of the current study. The current findings suggest that PrNMI is a promising opioid alternative for pain relief in the context of neuropathic pain.

## Author Contributions

AS, LML, EE, CC, and IS contributed to the design and writing of the manuscript. AS and DA contributed to the chronic pain behavior measurements. EE performed the conditioned place preference experiments and LML performed the intravenous self-administration experiments. CC and IS procured funding for the study.

## Statements and Declarations

### Ethical considerations

Procedures were conducted in accordance with the National Institutes of Health Guide for the Care and Use of Laboratory Animals, approved by the University of California Los Angeles Institutional Animal Care and Use Committee prior to experimentation, and were compliant with ARRIVE 2.0 guidelines.

### Declaration of conflicting interest

The author(s) declared no potential conflicts of interest with respect to the research, authorship, and/or publication of this article.

### Funding Statement

Research reported in this publication was supported by NIH grants CA196263, UL1TR001881, UG3NS128148 and the Shirley and Stefan Hatos Foundation.

### Data availability

The datasets generated during the current study are available from the corresponding author on request.

### Affiliation change

Amie Severino has since moved to Mount St. Mary’s University.

## Acknowledgements

We thank Herbert H. Seltzman (RTI International, NC) for the gift of PrNMI.

## Notes

### Competing Interest Statement

The authors have declared no competing interest.

